# Low-level, prediction-based sensory and motor processes are unimpaired in Autism

**DOI:** 10.1101/2020.09.01.277160

**Authors:** Johanna Finnemann, Kate Plaisted-Grant, James Moore, Christoph Teufel, Paul Fletcher

## Abstract

A new promising account of human brain function suggests that sensory cortices try to optimise information processing via predictions that are based on prior experiences. The brain is thus likened to a probabilistic prediction machine. There has been a growing – though inconsistent – literature to suggest that features of autism spectrum conditions (ASCs) are associated with a deficit in modelling the world through such prediction-based inference. However empirical evidence for differences in low-level sensorimotor predictions in autism is still lacking. One approach to examining predictive processing in the sensorimotor domain is in the context of self-generated (predictable) as opposed to externally-generated (less predictable) effects. We employed two complementary tasks – force-matching and intentional binding – which examine self-versus externally-generated action effects in terms of sensory attenuation and attentional binding respectively in adults with and without autism. The results show that autism was associated with normal levels of sensory attenuation of internally-generated force and with unaltered temporal attraction of voluntary actions and their outcomes. Thus, our results do not support a general deficit in predictive processing in autism.

## 1. Introduction

The predictive processing framework accounts for how we deal optimally with ambiguous signals from our environment using prediction-based optimisation of inference (Teufel and Fletcher [1], Friston and Kiebel [2]). While initially developed as a framework to understand healthy brain function, this account also offers potential insights into the processes underlying psychiatric disorders (Moore [3], Adams et al. [4], Barrett et al. [5], Sterzer et al. [6], Gadsby and Hohwy [7], Teufel and Fletcher [8], Corlett and Fletcher [9], Friston et al. [10], Kube et al. [11, 12], Fineberg et al. [13]). There has been a growing interest in applying this framework to investigate differences in the cognitive, perceptual and neural processes in autism spectrum conditions (Qian and Lipkin [14], Pellicano and Burr [15], Sinha et al. [16], Lawson et al. [17], Van de Cruys et al. [18], Rosenberg et al. [19], van Boxtel and Lu [20]). Much interest has been sparked by a proposal from Pellicano and Burr [15] suggesting that predictive deficits in individuals with autism are due to a diminished effect of prior expectations on the processing of ambiguous sensory information, leading to inferences that are more strongly based on sensory information. This atypicality in information processing, they speculate, could be a consequence of excessive endogenous neural noise although others have pointed out that reduced endogenous noise could yield comparable outcomes (Brock [21]). Alternative accounts suggest that the problem lies not in the prior expectations themselves but in altered precision of the prediction error – a key feedforward signal in the processing hierarchy (Van de Cruys et al. [22], Lawson et al. [17]).

Prima facie, the framework contributes a lot to understanding the characteristic clinical features of autism. For instance, it seems plausible to conjecture that deficits with the generation of predictions are at the core of difficulties with adapting to change, intolerance of uncertainty and certain sensory atypicalities in individuals with autism. Empirically, the evidence for these theories is still sparse and the idea of a global “predictive impairment […] shared across individuals” (Sinha et al. [16]) seems to be contradicted by an absence of apparent deficits in motion prediction of objects (Tewolde et al. [23]), predictions about the weight of objects based on material cues (Arthur et al. [24]) and other cognitive processes supposed to tap into predictive abilities (Croydon et al. [25], Manning et al. [26], Cruys et al. [27], Maule et al. [28]). Where group differences have been found, they mostly pertain to predictive deficits in the social domain: Balsters et al. [29], Chambon et al. [30], Turi et al. [31], Amoruso et al. [32], von der Lühe et al. [33], but this is not universally true, as Pell and colleagues have found no deficits in prediction-based perception of other people’s gaze direction (Pell et al. [34]). It is also unclear whether the observed deficits in prediction are due to low-level atypicalities in the predictive architecture or whether they might be the result of differences in other areas that prediction taps into such as the learning of action-outcome contingencies (Schuwerk et al. [35]) and temporal processing (Brodeur et al. [36], Szelag et al. [37]).

In short, while a predictive processing deficit provides a credible explanatory model for features of autism, the experimental evidence is currently inconsistent and requires clarification. Moreover, all of the paradigms mentioned above tap into higher-order perceptual and cognitive functions. In order to support the idea of a global prediction deficit in autism, however, a characterisation of basic mechanisms of sensory and motor prediction are currently lacking. These basic predictive mechanisms initially laid the foundations for the predictive processing framework (Holst and Mittelstaedt [38], Helmholtz [39]) but, surprisingly, have not been studied in ASD. In the current study we therefore used two complementary tasks known to index predictive processing in basic sensory and motor function: the forcematching task (Shergill et al. [40]) and a modified version of the intentional binding paradigm (Moore and Haggard [41]). We chose these tasks for two reasons: Firstly, in contrast to the higher-order cognitive paradigms mentioned above, both experiments focus on basic mechanisms of sensory and motor prediction that laid the foundations for the predictive processing framework ([38]). Secondly the tasks have robustly and reliably elicited responses in line with current views on prediction in healthy individuals and have, moreover, established the presence of altered responses in populations whose predictive architecture is conjectured to be compromised (Shergill et al. [42], Voss et al. [43], Synofzik et al. [44]).

The forcematching task measures attenuation of the sensory consequences of self-generated actions. It is based on the principle of motor control theory which suggests that sensory consequences of predictable forces are anticipated and attenuated. Tasks exploring this phenomenon have reliably demonstrated that self-generated sensory consequences are perceived as weaker than externally-generated sensory consequences of the same intensity across a range of experimental paradigms, volunteers and laboratories (Wolpe et al. [45, 46], Shergill et al. [40, 42], Voss et al. [47], Teufel et al. [48], Walsh et al. [49], Therrien et al. [50], Pareés et al. [51]).

The intentional binding (IB) effect refers to the finding that self-generated, voluntary actions and their sensory consequences are perceived to be closer together in time than movements externally forced upon the person and their sensory outcomes (Haggard et al. [52], Prinz and Hommel [53]). IB is thought to be an implicit measure of sense of agency (SoA) which in contrast to the sensory attenuation observed in the forcematching task, is speculated to rely both on predictive mechanisms as well as post-hoc inferences. Predictive and postdictive contributions to agency have been investigated by varying the probability with which the voluntary action produces the sensory out-come (Moore and Haggard [41]). Moore and Haggard found that both processes operate, but that one dominates depending on the specific outcome probabilities: On trials, on which the action produced an outcome with a high probability, healthy volunteers exhibited temporal binding even in the absence of the outcome, whereas subjective temporal compression was only observed on those low “outcome probability” trials that did indeed produce the outcome.

Thus, these two complementary tasks are well-suited to exploring different aspects of the predictive processing model of ASC: While the forcematching task is more likely to tap into basic predictive mechanisms of sensory gating (Chapman and Beauchamp [54], Hughes et al. [55]), intentional binding is thought to be largely attributable to temporal control and prediction (of the timing of the outcome). Therefore unimpaired performance on one, but not the other task would yield additional insight as to whether differences in predictive abilities in autism are more likely due to primary sensory deficits or more general issues with the timing and learning of action-outcome contingencies.

## 2. Experiment 1 – Forcematching in Autism

### 2.1. Method

#### 2.1.1. Participants

27 volunteers with a clinical diagnosis of an autism spectrum disorder and 26 healthy control participants (with no history of neurological or psychiatric illness) took part in the study. Written informed consent was obtained from all participants. Cognitive function for all study volunteers was assessed using the timed version of the Ravens Advanced Progressive Matrices (RAPM) (Raven et al. [56]) and the Wechsler FSIQ in the case of one ASC volunteer. Furthermore all participants filled in the Edinburgh Handedness Inventory [57] as handedness can have an effect on force-perception and production (Park et al. [58], Gertz et al. [59]). On the inventory, a score of +40 reflects right-handedness and a score below −40 left-handedness.

3 ASC participants were excluded from the subsequent analysis as two had a diagnosis of schizophrenia or another psychotic disorder and one was unable to complete the experiment due to difficulties with maintaining the required arm posture. Aside from psychotic disorders no other psychiatric conditions served as exclusion criteria as anxiety, depression, OCD and other neurodevelopmental disorders such as ADHD and dyspraxia are thought to be extremely common/co-morbid in ASC (for prevalence estimates see Leyfer et al. [60], Eaves and Ho [61], White et al. [62]). 10 of the participants with autism had co-morbid diagnoses of depression and/or anxiety and 6 were currently taking SSRIs. A further two people had a diagnosis of ADHD (one on medication) and one had unmedicated OCD.

Participants were well-matched for age, IQ (IQ information was unavailable for one control participant) and gender but the groups differed on the Edinburgh Handedness Inventory with three left-handed volunteers in the ASC group and none in the controls (see Table 1).

**Table 1:**
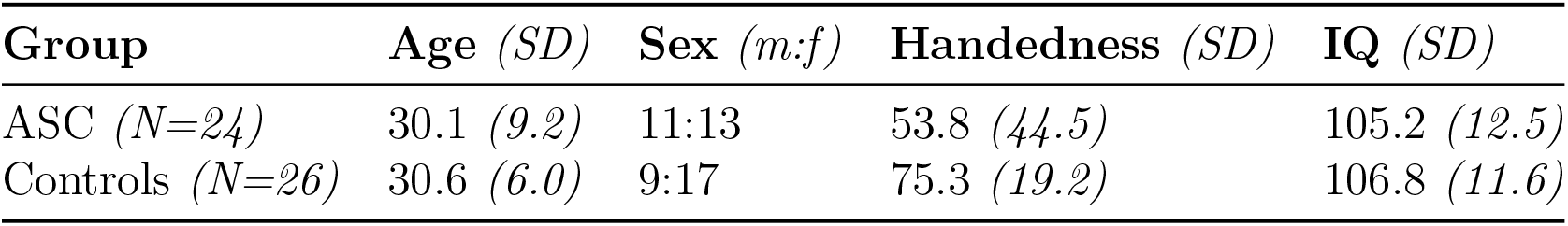
Participant Demographics

| Group | Age (*SD*) | Sex (*m:f*) | Handedness (*SD*) | IQ (*SD*) |
| --- | --- | --- | --- | --- |
| ASC (*N=24*) | 30.1 (*9.2*) | 11:13 | 53.8 (*44.5*) | 105.2 (*12.5*) |
| Controls (*N=26*) | 30.6 (*6.0*) | 9:17 | 75.3 (*19.2*) | 106.8 (*11.6*) |

All but 3 of the ASC participants were assessed with module 4 of the Autism Diagnostic Observation Schedule (ADOS, [63]) and while the group was moderately symptomatic (mean score: 6.7), only 9 participants met cut-off criteria for an autism spectrum condition and none met diagnostic criteria for autism. Low sensitivity of the ADOS module 4 has previously been reported and attributed to compensatory behaviour and “milder ASDs” ([64]). Even among children, those with a diagnosis of an autism spectrum condition that is not “childhood autism” (ICD-10) often do not meet the diagnostic cut-off for the ADOS (Baird et al. [65]).

Given previous reports of altered forcematching in individuals with high levels of schizotypy (Teufel et al. [48]), we used the 21-item Peters Delusion Inventory (PDI, Peters and Garety [66]) to quantify schizotypal traits in all participants. The Autism Spectrum Quotient (AQ, Baron-Cohen et al. [67]), a 50-item self-administered questionnaire, was used as a measure of autistic traits. AQ and PDI scores were unavailable for one ASC participant.

#### 2.1.2. Experimental Procedure

The experiment was modelled on the design by Shergill et al. [40] in which a lever – via a torque motor – exerts mild pressure onto the participants’ left index finger. Depending on the condition, participants were asked to match the experienced pressure to the point of subjective equality (i.e. the point where the pressure felt the same) by either pressing directly on the lever with their right index finger (“finger condition”) or by adjusting a slider which controlled the torque motor (“slider condition”), see Figure 1.

**Figure 1:**
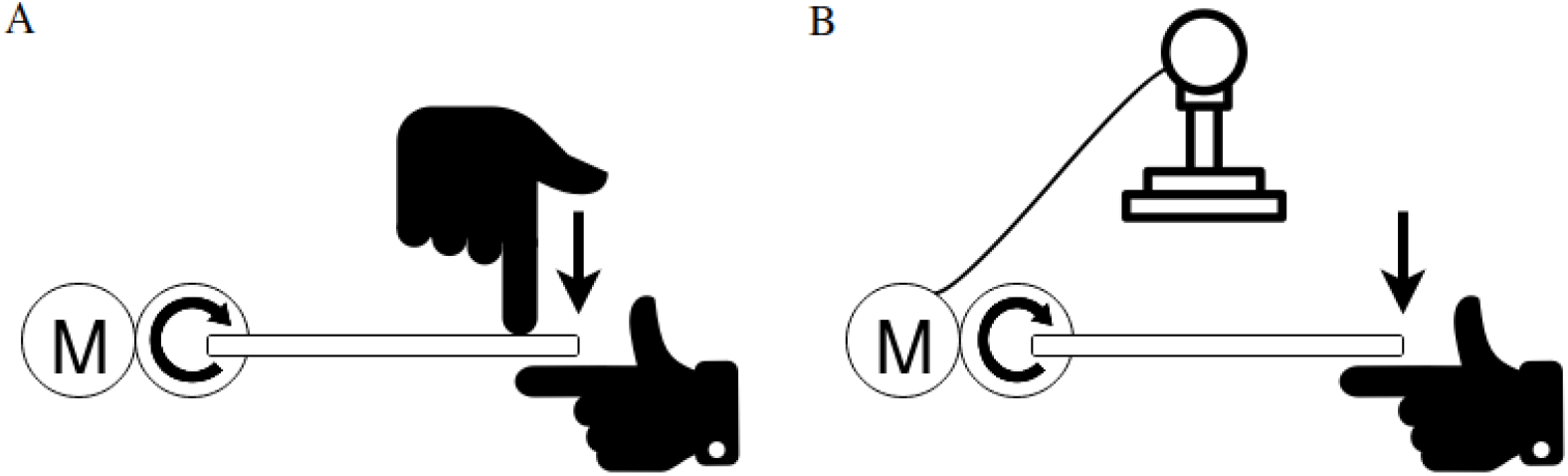
Illustration of the forcematching paradigm in which participants are asked to match a force applied to their left index finger via a lever. Participants had to reproduce the experienced force either by pushing down on the lever with their other index finger **(A)** or by moving a slider **(B)**.

As a result of forward prediction models for self-generated movements, participants routinely exceed the target force in the “finger” condition due to sensory attenuation, whereas predictions for the indirect control of the lever via the slider are less precise and participants thus tend to be more accurate in their reproduction of the force.

The slider was a potentiometer which transduced a force gain at the ratio of 0.5 N/cm. The target force was presented for 2.5 seconds (ramped up and down linearly over 0.25 seconds) after which an auditory go-signal indicated that participants should make their response to ensure that the matching took place within 2 seconds of the target force being withdrawn. After 3 seconds a second auditory signal indicated the end of each trial and instructed participants to lift their right index finger from the lever or move the slider back to the starting position. Mean force production was measured between 2 and 2.5 seconds after the start of the matching period, as in previous studies (Voss et al. [47]). Within each condition 10 different force magnitudes between 0.5N and 2.75N, differing in steps of 0.25N were applied in randomised order. Each force magnitude was presented for a total of 8 trials. Subjects first completed a 5-trial practice session for both conditions to ensure that they understood the task and were able to respond within the required time window. They then completed one “finger” and one “slider” block with 80 trials (160 trials in total). Invalid trials due to too slow or fast responses were repeated until a total of 80 valid trials had been completed. Practice sessions and test blocks were counterbalanced across both experimental groups.

#### 2.1.3. Data Analysis

One ASC participant was excluded from further analysis as their performance in the “finger” condition was more than 9 standard deviations above the mean.

Basic force attenuation was indexed by calculating an overcompensation score based on the difference between the matched forces in the “finger” and “slider” condition (each normalised against the passively experienced force) for each force level (see Humpston et al. [68]). Individual regression lines of target force versus matched force for each subject were fitted for the “finger” and “slider” condition and then summarised as group regressions for both conditions. In addition to the basic overcompensation score, the slope and intercept of the regression lines can provide more detailed information about the matching performance of different groups (Wolpe et al. [45]).

Group differences were evaluated with Bayesian estimation using Markov Chain Monte Carlo methods to generate samples of the relevant posterior distributions. JAGS (Plummer [69]) was implemented to build a Gibbs sampler and the default non-informative priors of the R package *BEST* (Kruschke [70]) were implemented. The data is assumed to follow a t-distribution in *BEST* with *v* (1-*∞*) degrees of freedom controlling the width of the tails and thus acting as a measure of normality. The wide priors make the estimation of the posterior parameters (mean(s) *µ*, standard deviation(s) *σ* and the shared normality parameter *v*) very data driven. Convergence was assumed as long as the Brooks-Gelman-Rubin scale reduction factor (Gelman and Rubin [71], Brooks and Gelman [72]) was *<*1.1. Bayesian correlations were calculated using the *BayesianFirstAid* package in R.

## 3. Results

Both groups showed the characteristic force attenuation with the posterior estimates of the mean overcompensation scores being 0.73 (credible interval/CI: [0.51, 1.00], estimated effect size: 1.58) and 0.80 (CI: [0.52, 1.10], estimated effect size: 1.33) for the control and autism group respectively. Handedness was unlikely to be associated with the magnitude of sensory attenuation (as measured by the overcompensation score) with an estimated correlation of *r*=−0.16 and a 95% CI of [−0.45, 0.16].

Plotting the mean linear regressions for matched forces in the “finger” and “slider” conditions did not suggest any group differences (Figure 2a). Congruously, Bayesian estimation yielded little evidence for a group difference on the means of overcompensation scores (estimated difference of means: −0.03, CI: [−0.37, 0.31], estimated effect size: −0.08, Figure 2b) or intercept (estimated difference of means: −0.04, CI: [−0.39, 031], estimated effect size: −0.09, Figure 2c) of the “finger” condition.

**Figure 2:**
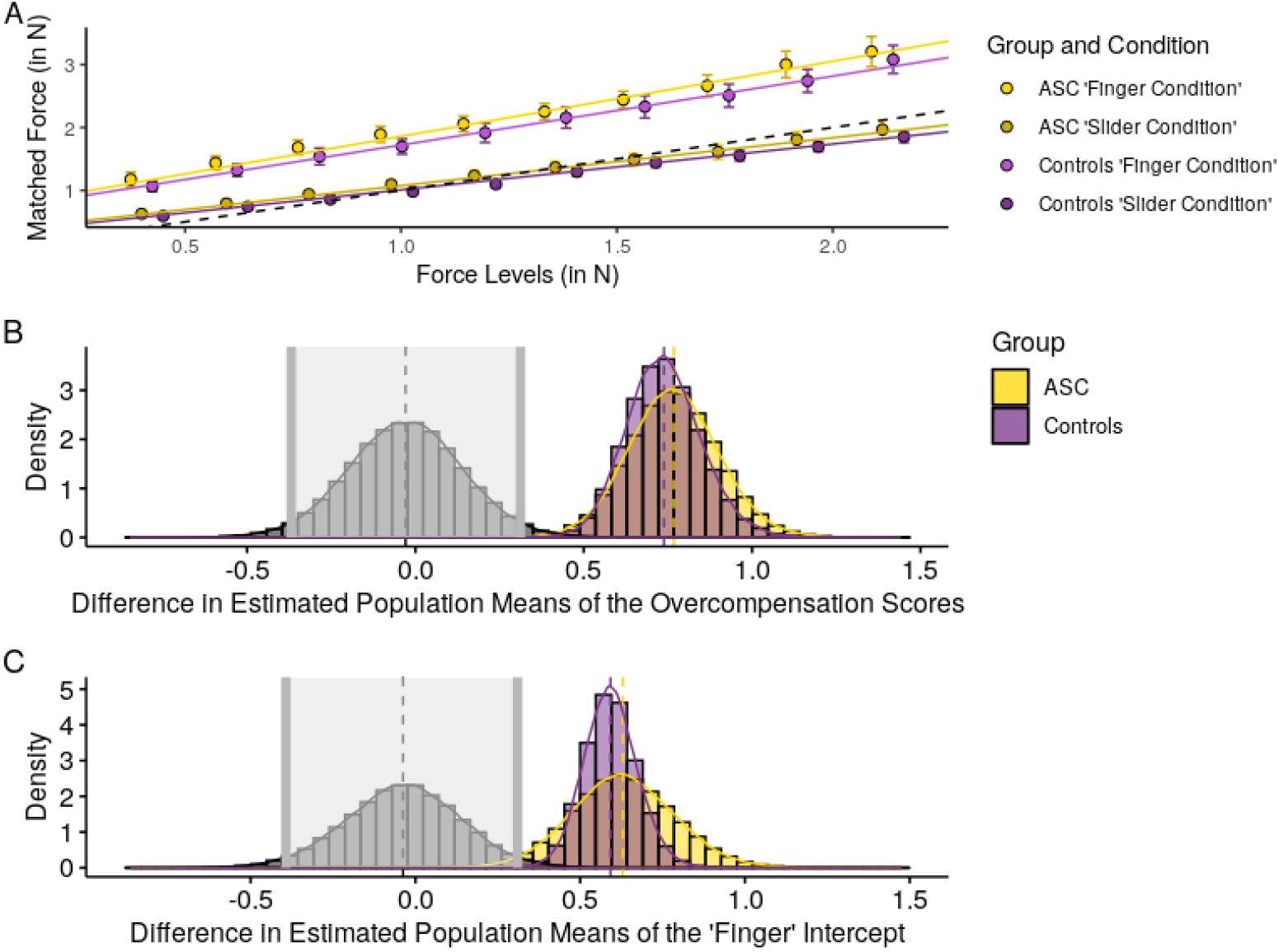
Main results for the forcematching task. **(A)** Mean linear regressions for the matched forces in the “finger” and “slider” conditions. Jitter was added to prevent overplotting. Error bars represent ±1 standard error (SE) of the mean. Perfect matching performance is indicated by the dashed black line. **(B)** A plot of the posterior probability of the difference in means for the overcompensation score (black) with the estimated population means in yellow and purple respectively. The shaded area is the credible interval (CI), in this case the 95% Highest Density Interval (HDI) **(C)** Posterior probability of the difference in means for the intercept in the “finger” condition.

For a more in-depth view at these measures see Appendix A.

### 3.0.1. Relationship between the Questionnaire Measures and Sensory Attenuation

As expected, posterior estimates for group means on the AQ indicated a difference (estimated difference of means: −19.49, CI: [−24.03, −15.06], estimated effect size: −2.62) and perhaps more surprisingly there was also evidence in favour of the true difference in means on the PDI being non-zero (estimated difference of means: −21.50, CI: [−42.22, −0.58], estimated effect size: −0.65) (Figure 3a).

**Figure 3:**
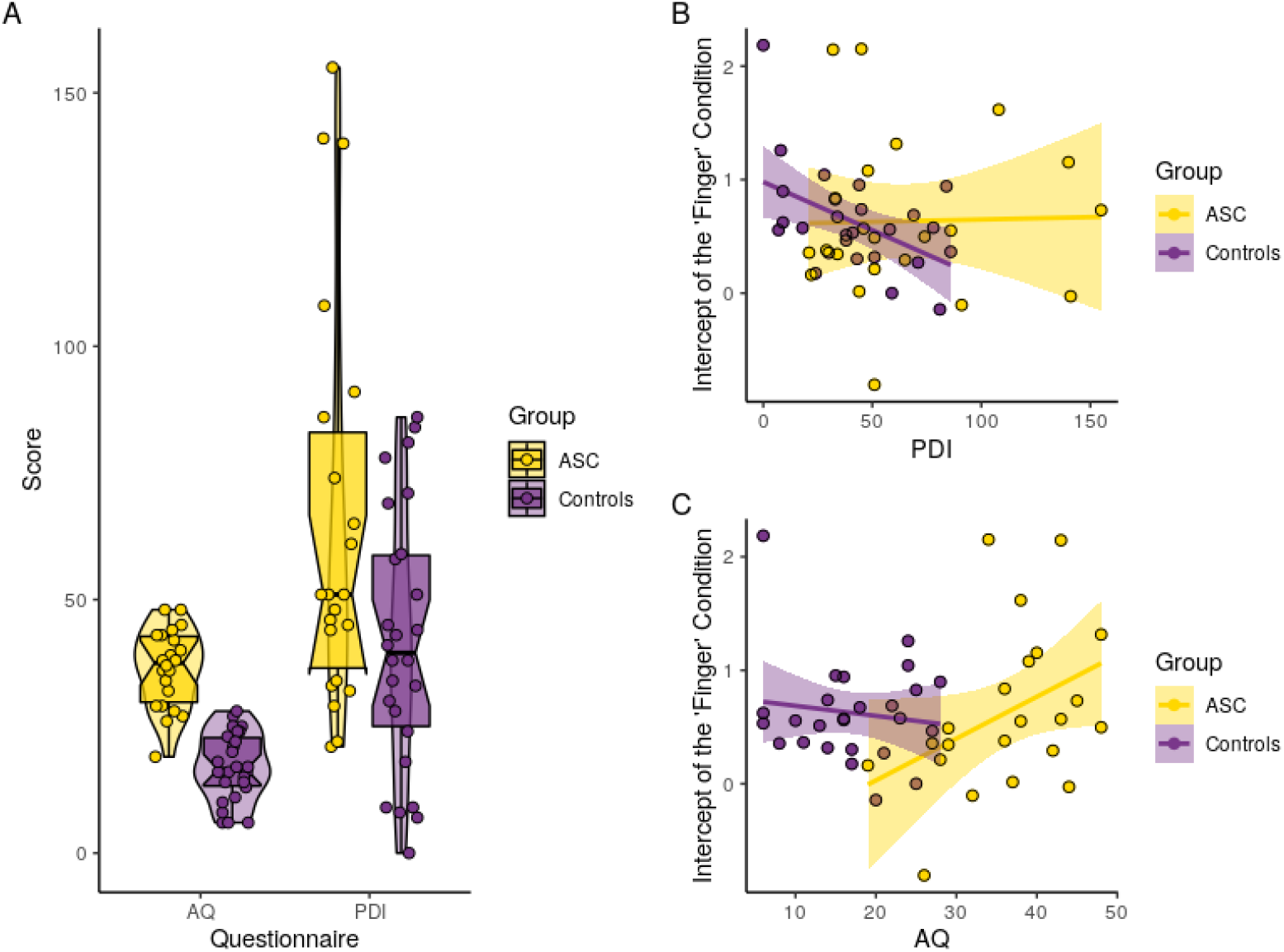
Results for the Questionnaire measures. **(A)** Plot showing the distribution of the questionnaire scores for both groups, including the median and interquartile ranges. **(B)** The correlation between sensory attenuation (as measured by the intercept in the “finger” condition) and the PDI. **(C)** The correlation between sensory attenuation (as measured by the intercept in the “finger” condition) and the AQ.

Using the intercept in the internal condition as the main measure of sensory attenuation (see: Wolpe et al. [45]), in line with previous observations (Teufel et al. [48]; but see: Humpston et al. [68]), we found that the probability that sensory attenuation has a negative relationship with schizotypy in the control group (probability: 98%, estimated correlation: −0.41, CI: [−0.73, −0.07]), whereas evidence in the ASC group suggested no significant relationship (estimated correlation: 0.04, CI: [−0.40, 0.45]). Conversely there did not seem to be an association between self-reported autistic traits on the AQ and sensory attenuation in the control group (estimated correlation: −0.01, CI: [0.42, 0.40]), but a trend for a positive relationship in the ASC group (estimated correlation: 0.36, CI:[−0.03, 0.70]), see Figure 3b and 3c.

### 3.0.2. Summary

Overall, we found no evidence of a deficit in the attenuation of self-produced sensory consequences in autism, which is in contradiction of existing predictive processing models of the condition. A Bayesian analysis supported an absence of group differences in key measures of sensory attenuation. Interestingly, not only AQ (as predicted) but also a measure related to schizotypy (PDI) was higher in the ASC group. Moreover, in line with previous work, correlative analyses of sensory attenuation with schizotypy showed an expected negative relationship in control participants. No such correlation was found in ASC. Conversely, AQ scores in the autism group correlated positively with sensory attenuation.

## 4. Experiment 2 – Intentional Binding in Autism

### 4.1. Method

#### 4.1.1. Participants

A total of 50 participants (25 per group) were recruited for the study. Written informed consent was obtained from all participants. All but one of the ASC volunteers also took part in experiment 1 and thus the same two volunteers with a history of psychosis were excluded.

Participants were matched for age, IQ (IQ information was unavailable for two control participants) and gender (see Table 2).

**Table 2:** Participant Demographics for the Intentional Binding Task

| Group | Age (*SD*) | Sex (*m:f*) | IQ (*SD*) |
| --- | --- | --- | --- |
| ASC (*N=23*) | 29.0 (*6.1*) | 11:12 | 105.2 (*12.7*) |
| Controls (*N=25*) | 31.2 (*5.7*) | 10:15 | 104.6 (*10.6*) |

#### 4.1.2. Experimental Procedure

The basic structure of the task was similar to other intentional binding experiments (Haggard et al. [52]): Participants were instructed to press a key with their right index finger at a time of their own choosing which caused a tone 250ms later. While they were engaged in this task, a Libet clock (Libet et al. [73]) was visible in the middle of the screen with a clock-hand rotating at a rate of 2560ms per revolution. After the keypress, the clock-hand continued to rotate for a random amount of time. Participants were told to avoid pressing at “premeditated” clock positions.

In the “action block” condition, participants had to recall the time at which they pressed the key (i.e. recall where the clock-hand was pointing to when they performed the keypress) while in “tone blocks” participants were asked to enter the the clock-hand’s position when they heard the tone. As in Moore’s adapted version (Moore and Haggard [41]), the probability of the tone occurring was manipulated: In half of the blocks (2 per condition) the tone followed the key press 50% of the time while in the other half it happened 75% of the time (see Figure 4). When no tone occurred, participants were asked to report a dummy value. Participants were informed of the response requirement (time estimation of the key press or tone occurrence) immediately prior to the blocks which otherwise did not differ visually from each other. The order of blocks was randomised for each participant.

**Figure 4:**
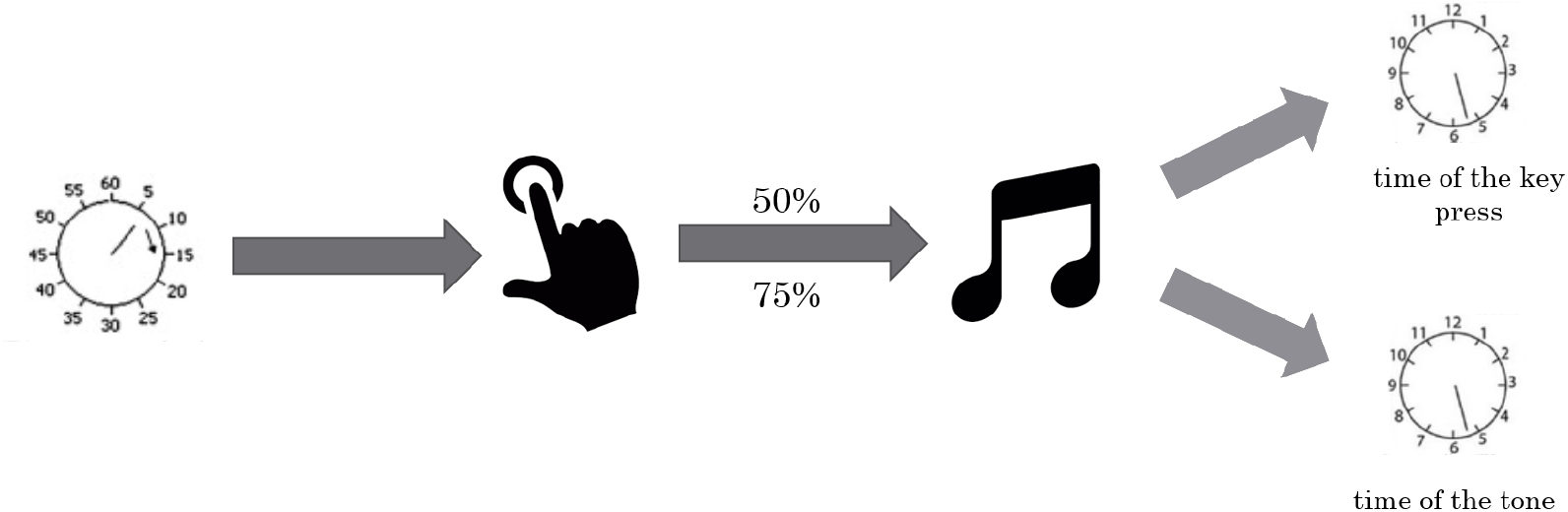
An illustration of the experimental procedure for IB with varying outcome contingencies

In addition to 8 experimental blocks (4 per condition), the volunteers also completed a baseline task requiring them to judge the time of their key presses without any subsequent tone.

Blocks with the 50% probability for tone occurrence had 50 trials whereas blocks with tones occurring 75% of the time had 40 trials. Baseline blocks had 50 trials. Due to a technical error 2 control subjects had the trial numbers reversed and 3 controls and 7 ASC participants only completed 40 trials in the baseline task.

The data from one of the control participants was excluded prior to the analysis as it became clear in the debriefing that he had not been following the instructions.

#### 4.1.3. Data Analysis

The analysis followed the typical protocol for IB studies. Initially, responses were corrected against the mean of all baseline trials for each participant. For the purposes of the analysis, the first 10 trials of each block were not included as participants had to learn the contingencies. The reported shifts in the performed key presses were used as the measure of intentional binding. By convention, binding for actions is indicated by a positive difference.

Based on Voss et al. [43], the predictive component to the intentional binding effect was calculated as the difference in overall shift between action only trials in the high probability blocks and action only trials in the low probability blocks (“action only” trials (75%) – “action only” (50%)). Since the tone is observed in neither condition, any difference in the strength of binding must be due to the higher predictive power of the “action only” 75% probability blocks. Analogously the inferential contribution was defined as the average shift in “tone only” trials in the 50% blocks. The authors describe the 50% contingency as subjectively “random”, so participants should not be able to form helpful predictions. Therefore any binding effect must be due to an inferential component that acts on the temporal estimation process after the tone occurs.

## 5. Results

### 5.0.1. Basic Intentional Binding Effect

The resulting pattern resembled Moore and Haggard’s [41] results where intentional binding was observed in all conditions apart from the low-probability no-tone trials (see Figure 5).

**Figure 5:**
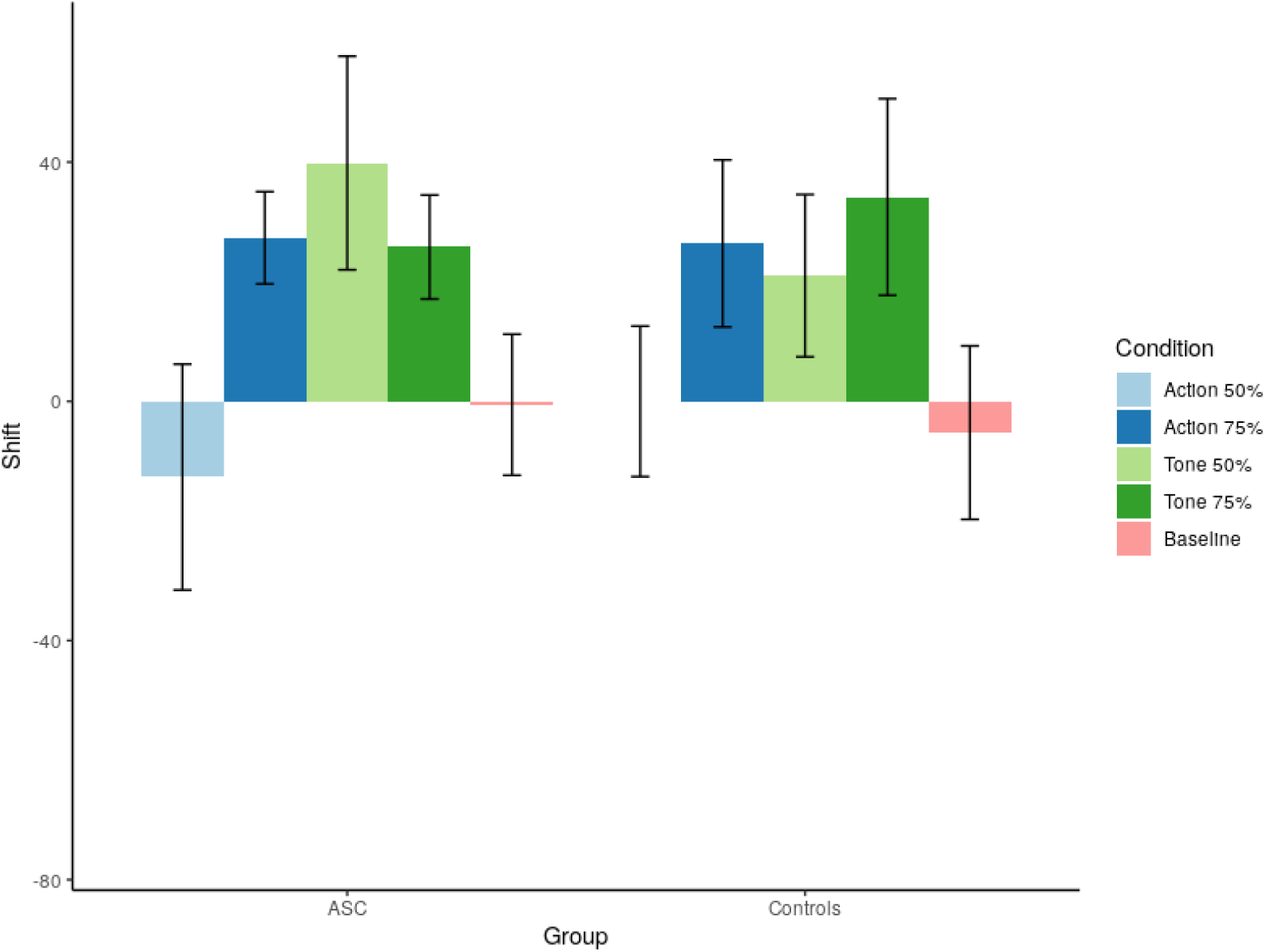
Baseline-corrected shift in the action estimates (ms) for each probability block in the “action only” and “tone only” conditions. Error bars represent ±1 standard error (SE) of the mean.

### 5.0.2. Group Comparison on Predictive and Inferential Components of Intentional Binding

The Bayesian estimation of the group difference for the predictive component (estimated difference of means: −13.7, CI: [−65.1, 37.9], estimated effect size: −0.17, Figure 6a) and the inferential component (estimated difference of means: −8.49, CI: [−59, 42.5], estimated effect size: −0.11, Figure 6b) makes a difference unlikely for both parameters.

**Figure 6:**
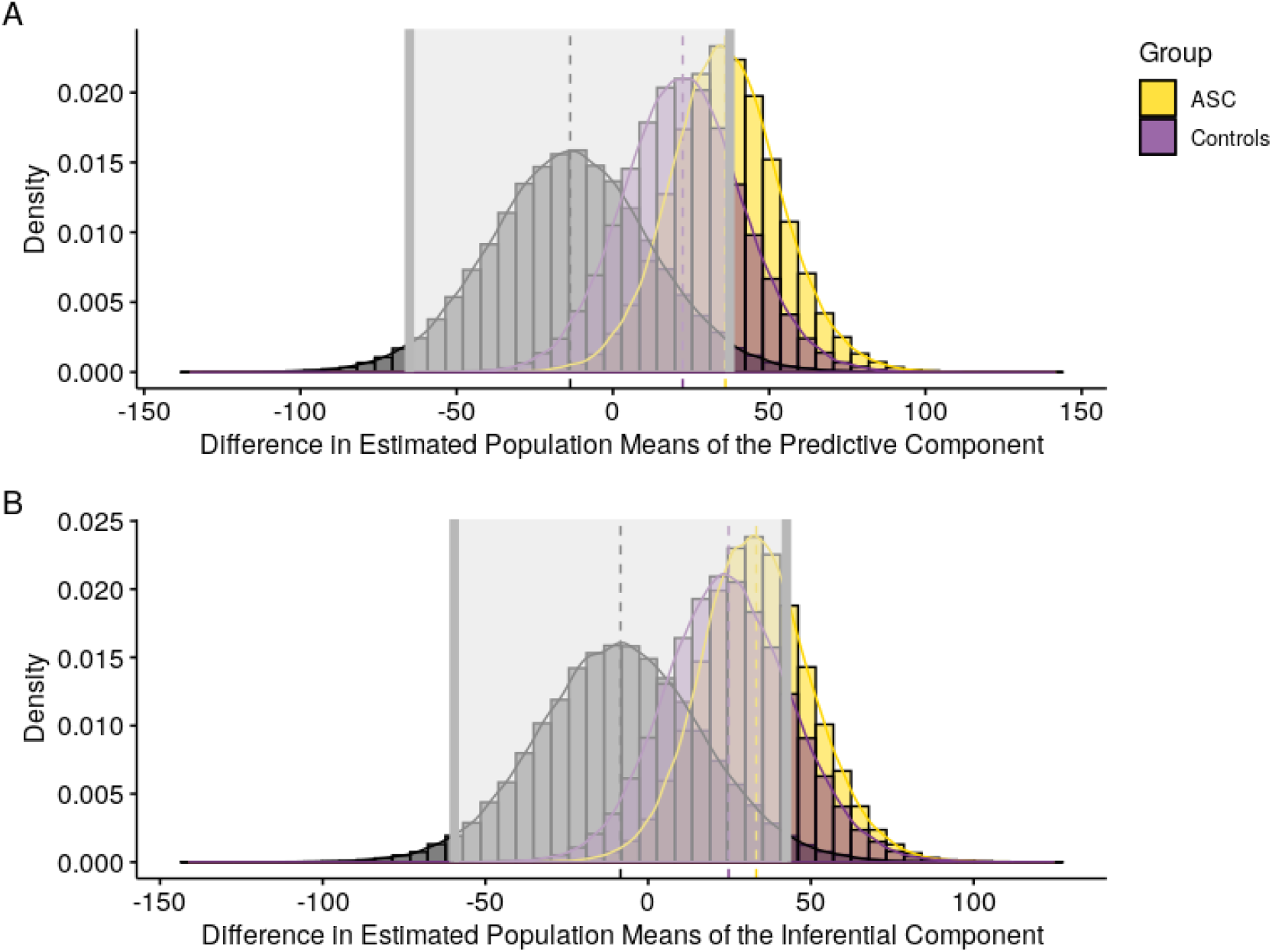
Posterior distributions for the difference in estimated population means of the predictive **(A)** and inferential **(B)** component of IB. The shaded area is the 95% Highest Density Interval (HDI).

### 5.0.3. Relationship between the Questionnaire Measures and Intentional Binding

There was little evidence that the AQ or PDI correlated with any of the measures; estimated correlations ranged between −0.22 and 0.23 and all CIs included 0.

### 5.0.4. Summary

Overall, therefore, in keeping with the findings from the force-matching task in experiment 1, we found no group difference in intentional binding. Both groups showed expected reductions in the subjective experience of action-outcome timing in both the predictive (tone absent) and postdictive (tone present) conditions.

## 6. Discussion

In the past decade, a number of prominent hypotheses have suggested that autism is primarily a disorder of atypical predictive processes and that the range of alterations, particularly in perceptual experiences can be explained in terms of these atypicalities. However the empirical evidence supporting these hypotheses in the form of differences in low-level sensorimotor prediction has been lacking which led us to investigate sensory attenuation and agency-based temporal binding in adults with autism. In light of this theoretical work conceptualising autism as a “disorder of prediction”(Sinha et al. [16]), one would expect to find reduced perceptual attenuation in the autistic group and a reduction of the predictive component to the intentional binding effect. Neither of these observations were made and our experiments do not support the idea of a deficit in predictive processing in autism. Both ASC and control groups demonstrated sensory attenuation of self-generated stimuli with a magnitude consistent with previously reported results (Teufel et al. [48], Shergill et al. [40], Wolpe et al. [45]) and both groups exhibited the basic pattern of inferential and predictive binding reported by Moore and Haggard [41]. These findings indicate that global deficits in predictive processing cannot explain the observed cognitive, perceptual and motor differences in autism spectrum conditions.

However, one interesting group difference that emerged lay in the within-group relationship between odd or unusual beliefs, as measured by PDI and the magnitude of sensory attenuation. While we replicated the previous finding that an increase in the number of delusion-like beliefs was associated with more accurate force-matching (i.e. reduced sensory attenuation), this relationship was not seen in autism. However there was some preliminary evidence that higher autistic traits in autistic individuals could be related to an increase in sensorimotor prediction as indicated by increased sensory attenuation. The lack of correlation between attenuation and PDI in the autism group is intriguing. One possibility is that the PDI and AQ questionnaires do not measure the same underlying traits in autism as in controls (Murray et al. [74]). An alternative explanation would be that sensory attenuation is indeed modulated by different latent traits in autistic and non-autistic individuals.

Compared to the schizophrenia literature, evidence for disruptions of sensory gating and agency processing in autism is scant: Previous research on sensory attenuation in ASC has reported unimpaired cancellation of self-generated tactile stimulation in the form of self-tickling (Blakemore et al. [75]) and adults with autism are just as good as their matched controls at judging agency based on whether visual feedback matched their own hand movements or not (David et al. [76]). In contrast, Zalla et al. [77] showed a decreased use of sensorimotor cues in making judgments of agency in adults with autism which was correlated with performance on a Theory of Mind task. They conclude that autistic individuals experience their internal signals as unreliable and might rely more on retrospective external cues (such as accuracy) to evaluate agency. Preliminary studies on interoceptive deficits in autism seem to support this claim (Noel et al. [78], Garfinkel et al. [79]). Similarly, Zalla and Sperduti [80] suggest that autism is characterised by an isolated impairment of predictive (but not postdictive) processes in the genesis of sense of agency. A recent study has indeed found an attenuated intentional binding effect in adults with autism when tested with visual, auditory and audio-visual action outcomes (Sperduti et al. [81]). In light of our diverging results the differences between the two experiments need to be examined: The manipulation of the probability of the action effect occurring in the experiment that is presented here is unlikely to cause an enhancement in overall IB, as it should introduce more uncertainty and more spurious binding effects. An obvious suggestion, given that Sperduti et al. employed three different delays between the action and action outcome, is that time estimation and temporal binding difficulties which are common in autism (Brock et al. [82], Maister and Plaisted-Grant [83]), impeded performance for the ASC group. As Maister and Plaisted-Grant [83] point out, impairments in estimating short time intervals between 0.5 and 2 seconds seem to be the result of deficits in attentional control in autistic individuals, rather than indicative of a more global temporal processing deficit and thus might elude being captured by the proportion error scores used in Sperduti et al. [81]. Other differences between the two studies include the smaller (N=15 for the autism group) all-male participant panel in Sperduti et al.’s experiment, the different estimation methods (Libet clock vs. analogue scale) and the fact that each condition (interval and modality) was only presented 10 times with 180 trials in total by Sperduti et al. compared to *∼*460 trials in the current study. If autistic individuals are indeed more variable in their responses due to attentional deficits, a higher number of trials would be needed to obtain the expected effect.

The lack of phenotyping for sensory reactivity and abnormalities is certainly a caveat of the present study and could be addressed more thoroughly in future investigations. Detailed assessments of sensory subtypes could also help to explain the commonly observed heterogeneity in task performance seen in the autistic group (Lane Alison E. et al. [84]) and it is possible that differences in predictive abilities might be domain-specific. As predictive attenuation is not unique to the tactile domain (Benazet et al. [85], Cardoso-Leite et al. [86], Desantis et al. [87], Hughes and Waszak [88]), an investigation linking domain-specific sensory reactivity (like the frequently reported auditory defensiveness) to sensory attenuation might be better equipped to uncover potential differences. Furthermore, although it is sometimes claimed that these sensorimotor processes are well understood given the extensive research into central and peripheral nervous system mechanisms supporting sensory gating (Rushton et al. [89]), their relationship with the perceptual attenuative processes seen in the force matching task is not entirely clear and there is some evidence that the two processes are functionally distinct (Palmer et al. [90]).

A further limitation of the experiments presented here was the exclusion of younger populations for the experiments. As autism is a neurodevelopmental disorder, it would be worth exploring if the trajectories for acquiring and refining internal models of the external world are different in autistic individuals even if performance is indistinguishable at a later developmental stage. Since structural priors are likely to either emerge from long-term aggregation of individual experiences or as embedded constraints acting on bottom-up processes (Teufel and Fletcher [1]) – as opposed to the short-term learning of stochastic relationships for contextual priors – they supposedly are subject to developmental processes. As such the force-matching task would be the best candidate for a developmental approach to predictive coding paradigms.

Our study aimed to explore the predictive abilities of individuals with autism in two motor tasks that are thought to be subserved by partially overlapping, but different neural mechanisms. Previous efforts to investigate predictive processing in autism have yielded inconclusive results (mostly supporting aberrant prediction in the social domain), despite a comparatively large theoretical literature. Our present study militates against the the idea of a general prediction deficit in autism as results indicate intact predictive and postdictive mechanisms of sensory attenuation and temporal attraction between actions and action outcomes. However results hinted at more subtle differences in the relationships between latent traits of schizotypy/autism and task performance in the two groups which illustrates the need to consider potential discrepancies in specific domains or subgroups.

