## Appendix for "Low-level, prediction-based sensory and motor processes are unimpaired in Autism"

### Appendices

#### A. Additional Analyses for the Forcematching Task

##### *A.1. Slope and Intercept Measures*

Bayesian estimation of the parameters for the slope of the regression for the "finger" condition did not indicate any group differences (estimated difference of means: -0.04, CI: [-0.35, 0.27], estimated effect size: -0.10).

For the "slider" condition, Bayesian estimation equally did not suggest any group differences for the slope (estimated difference of means: -0.01, CI: [-0.18, 0.16], estimated effect size: -0.04), which has been interpreted as a measure of sensory sensitivity (Wolpe et al. [45]).

##### *A.2. Noise and Precision*

In line with previous research (Wang et al. [91]), autistic participants demonstrated subtle motor deficits in force control as evidenced by noisier responses across both conditions: Mean-squared errors (MSE) for individual regression lines were computed (estimated difference of means: -0.18, CI: [-0.34, -0.02], estimated effect size: -0.78, Figure 7a).

One possible explanation for a worse regression fit in the autistic group may lie in the sampling method. If the volunteers with autism differed on parameters such as the overall time to stabilise (eg. with longer initial overshoots), downward drift (fatigue) or other factors affecting the overall shape of the force traces, the pre-determined time window might not be sampling the intended matched force accurately on every trial.

Thus, in an attempt to minimise the effect of the shape of individual force traces, an alternative measure was adapted from Wolpe et al. [45] which searched for the 0.5 second time window with the least amount of variability on each trial. The "time to stabilise", that is to say the time between the onset of movement and the calculated time window, did not differ between groups in either the "finger" (estimated difference of means: -32.2, CI: [-111, 44.2], estimated effect size: -0.287) or "slider" (estimated difference of means: -113, CI: [-245, 15.4], estimated effect size: -0.53) condition.

With this adjustment, no difference in the overcompensation scores could be detected (estimated difference of means: 0.054, CI: [-0.30, 0.42], estimated effect size: 0.34), see Figure 7b.

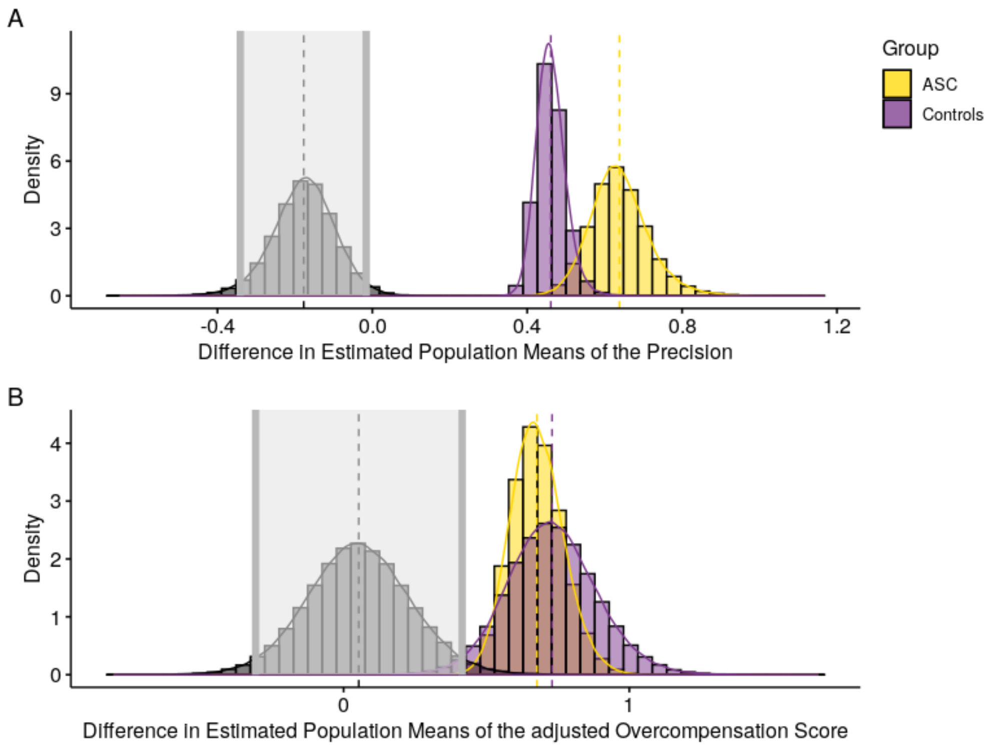


**Figure 7:** **(A)** Posterior probability distribution of the difference of means for individual MSEs. **(B)** Posterior probability distribution of the difference of means for the adjusted Overcompensation Score.
